# Topologically Dependent Abundance of Spontaneous DNA Damage in Single Human Cells

**DOI:** 10.1101/859686

**Authors:** Qiangyuan Zhu, Yichi Niu, Michael Gundry, Kuanwei Sheng, Muchun Niu, Chenghang Zong

## Abstract

In the studies of single-cell genomics, the large endeavor has been focused on the detection of the permanent changes in the genome. On the other hand, spontaneous DNA damage frequently occurs and results in transient single-stranded changes to the genome until they are repaired. So far, successful profiling of these dynamic changes has not been demonstrated by single-cell whole-genome amplification methods. Here we reported a novel single-cell WGA method: Linearly Produced Semiamplicon based Split Amplification Reaction (LPSSAR), which allows, for the first time, the genome-wide detection of the DNA damage associated single nucleotide variants (dSNVs) in single human cells. The sequence-based detection of dSNVs allows the direct characterization of the major damage signature that occurred in human cells. In the analysis of the abundance of dSNVs along the genome, we observed two modules of dSNV abundance, instead of a homogeneous abundance of dSNVs. Interestingly, we found that the two modules are associated with the A/B topological compartments of the genome. This result suggests that the genome topology directly influences genome stability. Furthermore, with the detection of a large number of dSNVs in single cells, we showed that only under a stringent filtering condition, can we distinguish the *de novo* mutations from the dSNVs and achieve a reliable estimation of the total level of *de novo* mutations in a single cell.

---

To study single-cell genomics, various single-cell whole genome amplification (WGA) methods have been developed to detect permanent genomic variations in single cells, including single nucleotide variations (SNVs) ^1-6^ and copy number variations (CNVs) ^7-9^. However, we have been ignoring the fact that spontaneous DNA damage frequently occurs to the genome, resulting in single-stranded changes ^10-13^. In comparison to *de novo* mutations, the measurement of single nucleotide damage including oxidation, deamination and alkylation etc. at single cell resolution is greatly desired for studying genome stability and the physiology of the cells. In theory, it is possible that we can detect DNA damage by single-cell whole genome amplification (WGA) methods since the damaged bases can lead to nucleotide misincorporation in amplification, which can then be detected as the potential *de novo* variants in the sequencing data. We denote these DNA damage associated single nucleotide variants as dSNVs.

So far, no single-cell WGA methods have been developed to detect these dynamic DNA damage. Two major technical obstacles in achieving reliable detection of *de novo* variants are amplification errors and technically introduced damage to DNA ^5,6^. Here we reported a new single-cell WGA method: Linearly Produced Semiamplicon based Split Amplification Reaction (LPSSAR). The reaction scheme of LPSSAR is shown in **Fig. 1**. In LPSSAR, after the cell is lysed at 30°C by an alkaline solution, the three annealing and extension cycle is performed (**Fig. 1**). In the three cycles, the semiamplicons are linearly copied from the genomic DNA. In contrast, the full amplicons are nonlinearly produced from the semiamplicons (**Fig. S1A**).

**Fig. 1:**
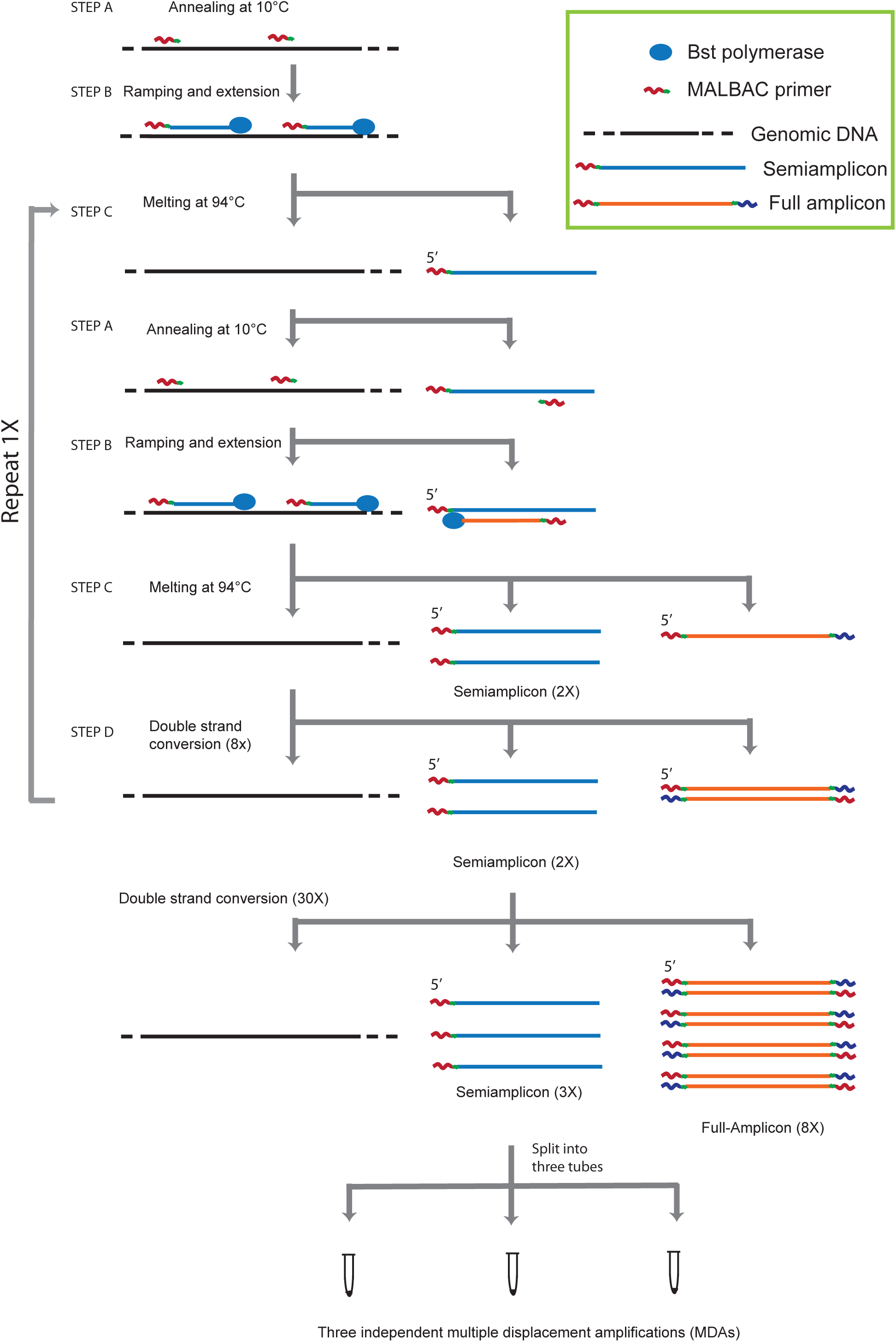
The reaction scheme of LPSSAR.

Next, we apply Multiple Displacement Amplification (MDA) for the second round of amplification. To warrant that only the linearly produced semiamplicons and original genomic DNA are amplified in the MDA, we first perform 30 cycles of PCR without the melting steps. In our diagram, we denote this step as double-strand conversion (DSC). As the result of DSC, the nonlinearly produced full amplicons are efficiently converted to double-strand DNA. Meanwhile, the semiamplicons remain as single-stranded DNA since they cannot hybridize with PCR primers. In the following MDA reactions, since the double-strand full amplicons cannot be primed by hexamer, therefore, only the linearly produced semiamplicons and original genomic DNA can be amplified. It is worth noting that if the full amplicons are used for amplification by PCR as illustrated in **Fig. 1**, it would correspond to the previously published MALBAC method by Zong *et al.* ^2^.

After DSC, the preamplification products are then split into three tubes for independent amplification by MDA (**Fig. 1**). A split-amplification scheme is necessary for effective filtering of the technical errors independently introduced during the amplification step and ensure accurate dSNV detection (**Fig. S1B)**. When we perform one MDA reaction in a single tube without splitting the preamplification products, amplification bias could lead to the overrepresentation of a small portion of semiamplicons. If amplification errors exist in these overamplified semiamplicons, these amplification errors will be called as *de novo* SNVs in the downstream analysis. In contrast, with the split-amplification strategy, we only call the variants that are detected in at least two independent reactions as *de novo* variants. Thus, the vast majority of the amplification errors are filtered out, therefore, ensuring the high accuracy in detecting biological *de novo* variants. This strategy requires robust MDA reactions of three splits, which is achieved by frequent mixing of the MDA reaction (**Fig. S2, Supplemental Materials**).

To evaluate the performance of LPSSAR, we amplified single cells isolated from a cultured cell line (MCF10A, a normal diploid breast epithelial cell line) and the single nuclei of human cortical neurons isolated from frozen brain samples. For MCF10A, three single cells were isolated from the serum-free culture in the poised G0 state to warrant the diploid genome. For human cortical neurons, we amplified and sequenced 18 single cells from three brain samples. Each cell was sequenced at 30X sequencing depth with ∼10X coverage for each split amplification. On average, we achieved genome coverage of ∼83% per split (**Table S1**).

Next, we combined the sequencing data of three splits and used GATK ^14^ for variant calling. Without the two-split detection criterion, we were able to estimate the amplification error rate to be ∼5×10^−5^ per base, consistent with the known error rate (4×10^−5^ per base) of Bst polymerase utilized in our preamplification reaction. The probability that two splits randomly introduce the same amplification error is (5×10^−5^)^2^×0.333≈8.3 ×10^−10^ per base (0.333 is the probability for the two splits to have exactly the same type of error), which corresponds to five false positives per genome caused by amplification errors, underscoring the successful filtering of the vast majority of amplification errors in our approach.

In **Fig. 2A**, we plotted the number of detected heterozygous germline mutations by allele frequency (AF) values (**Table S2**) for MCF10A cells. The Gaussian distribution centered at 0.5 demonstrates the evenness of our amplification. To identify *de novo* variants, we need to filter out all the germline SNVs detected in the bulk sequencing data. Here, we implemented the zero-read criterion: only the variants not detected in the bulk sequencing data were defined as *de novo* variants in single cells. The single-cell statistics of *de novo* variants can be found in **Table S3**. Surprisingly, even after effective filtering of amplification errors as described above, we still detected 10,634±1,845 *de novo* variants per MCF10A cell. In **Fig. 2B**, we plotted the distribution of AF values for these presumed *de novo* variants. These variants center at a low AF value (AF=0.1), in contrast to germline mutations which center at 0.5 (**Fig. 2A**). This different AF distribution strongly suggests that the detected *de novo* variants did not originate from mutations. Instead, the low AF value indicates that they are originated from single-stranded variants, which is likely to be associated with single-stranded DNA damage. The distribution of the AF values for single-stranded variants should center at 0.25, however, considering that the AF values of half of dSNVs are less than 0.25, we expect to observe that the AF distribution of dSNVs peaks at AF=0.1 as observed in **Fig. 2B**.

**Fig. 2:**
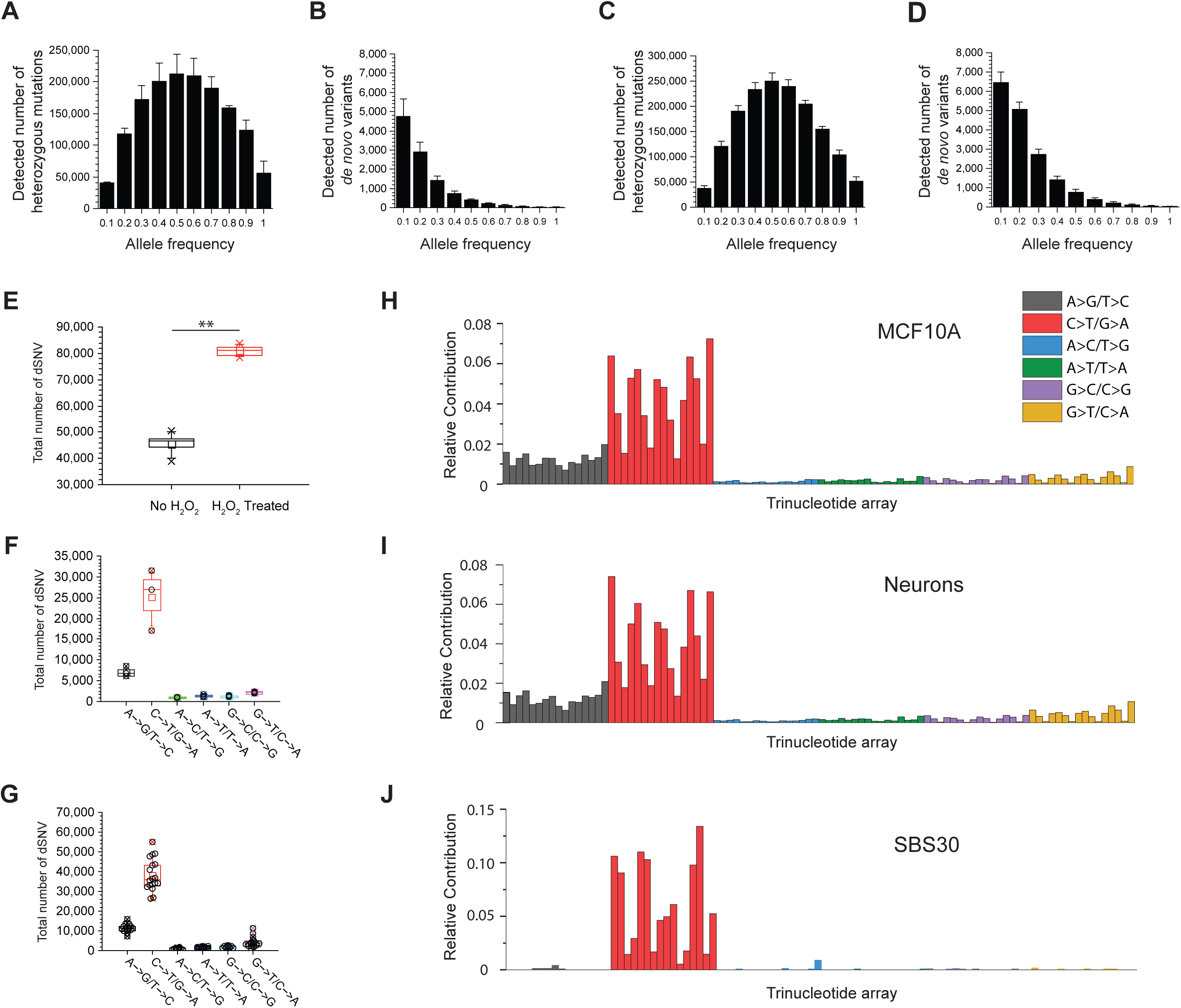
The measurement of dSNVs in MCF10A cells and neurons. (**A**) The histogram of allele frequency (AF) distribution of heterozygous germline mutations in single MCF10A cells. The Gaussian distribution centered at 0.5 indicates the evenness in our amplification. (**B**) The histogram of AF distribution of *de novo* variants in single MCF10A cells. The distribution centered at low AF values supports the detection of damage associated variants. (**C**) The histogram of AF distribution of heterozygous germline mutations in neurons. (**D**) The histogram of AF distribution of *de novo* variants in neurons. (**E**) The comparison of the total number of dSNVs between the MCF10A cells with and without hydrogen peroxide treatment (P < 0.01). (**F**) The total number of different types of dSNVs of the three single MCF10A cells at G0 phase. (**G**) The total number of different types of dSNVs of all sequenced neurons. (**H**) The trinucleotide signature of dSNVs detected in the MCF10A cells. The dSNV signature resembles COSMIC SBS30 (cosine similarity = 0.86). (**I**) The trinucleotide signature of dSNVs detected in all neurons. (**J**) The trinucleotide signature of COSMIC SBS30.

For single cortical neurons, we observed similar AF distributions for germline mutations and *de novo* variants (**Fig. 2C-D**). The average level of the detected dSNVs in 18 single neurons is 16,274±725 per neuron, which is significantly higher than the dSNVs detected in single MCF10A cells. Next, based on the genome coverage in each split, we can estimate the detection rate of dSNVs, and then the total level of dSNVs in single cells can be readily estimated (**Supplementary Materials** and **Table S3**).

To demonstrate the ability in detecting DNA damage, we applied LPSSAR to the cells damaged by hydrogen peroxide (H_2_O_2_). Only five minutes of H_2_O_2_ treatment was used to avoid introducing significant mutagenesis. After sequencing, we observed that the number of DNA damage is significantly increased in H_2_O_2_-treated cells (**Fig. 2E**). The increase of dSNVs in all six subtypes of variants is consistent with the known DNA lesion pattern caused by H_2_O_2_ treatment (**Fig. S3**) ^15^.

In **Fig. 2F-G**, we plotted the complete array of the detected dSNVs for MCF10A and cortical neurons respectively. Based on the knowledge of the major forms of DNA damage occurred to the genome, dSNVs of C->T/G->A are associated with cytosine oxidation ^16,17^, the dSNVs of A->G/T->C are potentially associated with adenine oxidation followed by deamination ^10^, and the dSNVs of G->T/C->A are associated with 8-oxoguanine ^18,19^. Among all six types of dSNVs, the major contribution to dSNVs is from the transition variants: C->T/G->A and A->G/T->C. Next, we compared the trinucleotide signature of dSNVs to the COSMIC mutational signatures ^20^. In **Fig. 2H-I**, we showed the trinucleotide signature of dSNVs for MCF10A cells and neurons and the similar signatures between MCF10A cells and neurons indicate the generic pattern of DNA damage exists across different cell types. The COSMIC mutational signature with the highest similarity score is SBS30 with cosine similarity of 0.86 (**Fig. 2J**). In a recent knockout study of *NTHL1* gene, SBS30 has been also shown as the mutational signature of *NTHL1* deficiency ^21^. Our result indicates that *NTHL1* gene encodes the proteins that repair the C->T/G->A damage --- the major component of dSNVs. Therefore, *NTHL1* deficiency could lead to severe mutagenesis. Indeed, it has been reported that *NTHL1* deficiency is highly mutagenic and promotes adenomatous polyposis and colorectal cancer ^22^.

Next, we calculated dSNV densities in different functional regions. In both MCF10A cells and neurons, different regions have different dSNV densities (**Fig. 3A-B, Table S4**), which strongly supports that the dSNVs detected by LPSSAR are biologically genuine. We observed that the dSNV densities in the exon and promoter regions are significantly lower than the intron and intergenic regions (p < 0.0001). Between exon and promoter regions, the dSNV densities in promoter regions are lower (p < 0.0001). Between the intron and intergenic regions, dSNV densities in the intergenic regions are lower (p = 0.018). The potential mechanisms for all these differences are likely due to different levels of DNA assault or different levels of repairing efficiency in different functional regions.

**Fig. 3:**
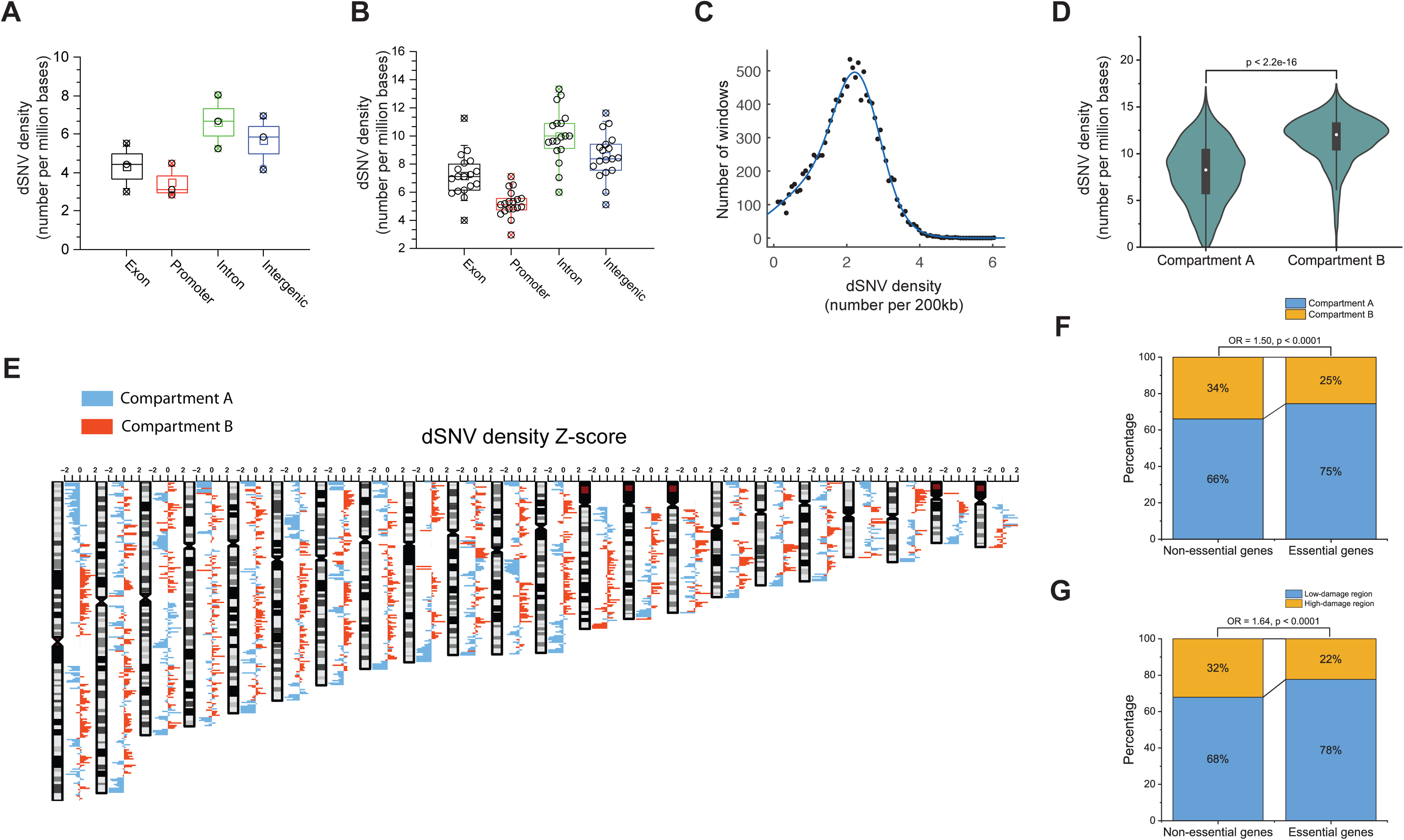
The genomic distribution of dSNVs in MCF10A cells and neurons. (**A**) The dSNV densities (number per million bases) in different genomic regions (exon, promoter, intron, and intergenic regions) in MCF10A cells. (**B**) The dSNV densities in different genomic regions (exon, promoter, intron, and intergenic regions) in all neurons. (**C**) The histogram of the dSNV densities in 200 kb binning windows in the genome of neurons. The distribution is fitted into two Gaussians with 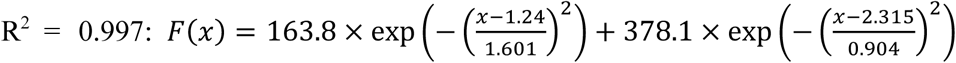 by least absolute residuals (LAR). (**D**) The dSNV densities of neurons in A compartments and B compartments. A/B compartment is determined at 1 Mb resolution. (**E**) The Z-score of the dSNV density in each binning window along the genome (window size: 1 Mb). The A compartments are blue colored, and B compartments are red colored. (**F**) Essential genes are more enriched in the A compartments than the B compartments. (**G**) Essential genes are more enriched in the low-damage regions (dSNV density Z-score < 0) than the high-damage regions (dSNV density Z-score > 0).

To further investigate the distribution of dSNVs on the genome of neurons, we plotted the histogram of the dSNV densities in 200 kb windows of genome (**Fig. 3C**). The total number of 18 neurons allows the sufficient sampling of dSNV density in each binning window. The distribution of dSNV densities can be fitted into two Gaussians with the goodness of R^2^=0.99 (**Fig. 3C**), indicating the existence of two types of DNA damage modules: high-damage and low-damage modules. In the topological profiling of the genome by Hi-C experiments, A/B compartments have been observed as a generic feature of 3D genome across different cell types ^23-27^. To investigate the potential relation between our DNA damage distribution and topological compartments in human prefrontal cortex ^24^, we compared the dSNV densities in the A compartment versus B compartment. As shown in **Fig. 3D**, the significant difference between the dSNV densities in the A and B compartments suggests that the abundance of DNA damage depends on the topological structure of the genome.

Next, we calculated the Z score of dSNV density for each window. In **Fig. 3E**, we plotted the Z scores of each window along the chromosome: the Z-score is shown as the height of the bar. The color indicates whether the window belongs to A or B compartment. We observed that the majority of the binning windows with positive Z-scores (high dSNV density) are red colored (B compartment); the majority of the binning windows with negative Z-scores (low dSNV density) are blue colored (A compartment). The consistent pattern between Z-score and the compartment assignment strongly indicates that A/B compartments influence the abundance of DNA damage along the genome. The reason for this phenomenon could be due to that DNA repair is less efficient due to limited accessibility to genome in the relatively closed B compartments. Furthermore, we showed that the human essential genes are preferentially located in A compartments (**Fig. 3F**) as well as in the low-damage regions (**Fig. 3G**). This observation indicates that the genome topology shapes the landscape of DNA damage along the genome, which results in different enrichment of essential genes in different topological compartments. The list of human essential genes was summarized based on previous studies ^28-31^.

Next, with the observation of the large number of dSNVs existing in single cells, it is important to point out that without an effective filtering of dSNVs from *de novo* mutations, we cannot achieve a reliable estimation of the total levels of *de novo* mutations in single cells. The underlying reason is that the amplification bias, which leads to different scenarios of strand dropouts as shown in **Fig. 4A**, makes it difficult to distinguish mutations from dSNVs. To test different criterions for detecting *de novo* mutations, we performed a single-cell expansion experiment as shown in **Fig. 4B**. Firstly, by comparing the sequencing data of Bulk S1 and S2, we identified the somatic mutations existing in the original cell selected for single-cell expansion (937 somatic mutations with TsTv ratio of 0.87). Next, we performed LPSSAR for three single cells isolated from the expanded clone. With the three single-cell sequencing data, different criterions for filtering out the dSNVs were then tested. For transition variants, we found out that under the following filtering criterion: the variants were detected in all three splits with AF ≥0.4 and Q (variant calling score) ≥ 800, the majority of dSNVs can be filtered out (**Fig. 5A**). For transversion variants, with the same AF criterion, the Q cutoff value can be reduced to 400 (**Fig. 5B**). In contrast, under the criterion of the detection in at least two splits, we cannot effectively separate *de novo* mutations from dSNVs (**Fig. S4A-B**). This underscores the necessity of using three splits for distinguishing the true *de novo* mutations from the dSNVs.

**Fig. 4:**
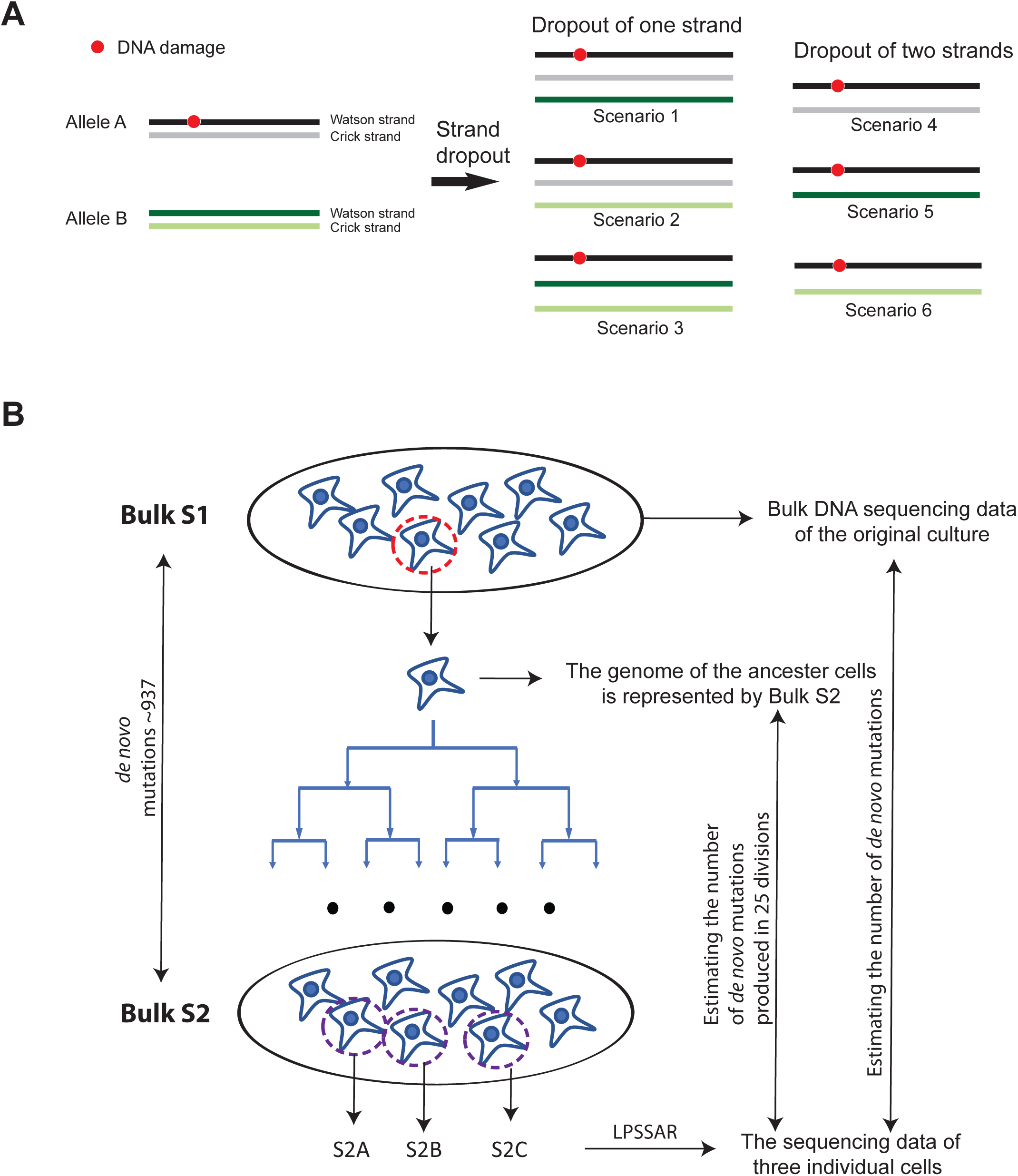
The diagrams for strand dropout and single-cell expansion. (**A**) Different scenarios of strand dropout causing the technical challenge in distinguishing *de novo* mutations from dSNVs. (**B**) The single-cell expansion scheme for validating the accuracy in estimating the total number of *de novo* mutations in single cells.

**Fig. 5:**
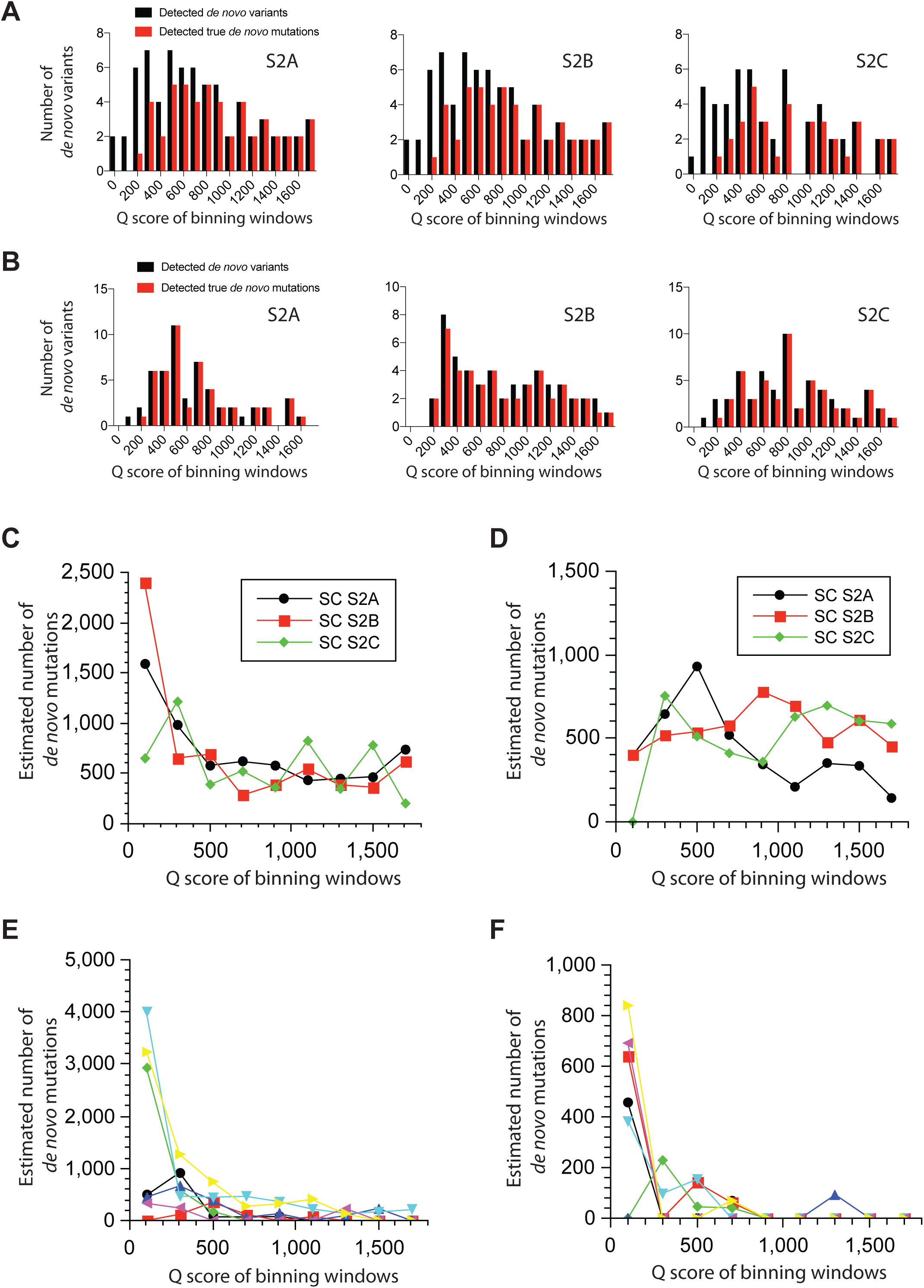
The estimation of the total number of *de novo* mutations in MCF10A cells. (**A**) The numbers of the detected *de novo* transition variants and the true *de novo* transition mutations in the binning windows of the variant calling scores (Q scores) with the AF>0.4 in all three splits criterion. (**B**) The numbers of the detected *de novo* transversion variants and the true *de novo* transversion mutations in the binning windows of the Q scores with the AF>0.4 in all three splits criterion. (**C**) The estimated total number of *de novo* transition mutations on the axis of the Q scores of the binning windows using the stringent AF criterion for the three cells isolated from the single-cell expanded culture. The reference bulk is Bulk S1. (**D**) The estimated total number of *de novo* transversion mutations on the axis of the Q scores of the binning windows using the stringent AF criterion for the three cells isolated from the single-cell expanded culture. The reference bulk is Bulk S1. (**E**) The estimated total number of *de novo* transition mutations on the axis of the Q scores of the binning windows using the stringent AF criterion for the neurons of the brain sample p5554. (**F**) The estimated total number of *de novo* transversion mutations on the axis of the Q score of the binning windows using the stringent AF criterion for the neurons of the brain sample p5554.

To estimate the total number of *de novo* mutations in single cells, we first determine the detected germline mutations in each binning window of Q values larger than the Q cutoff value derived from **Fig. 5A-B**. The detection rate of the germline under the stringent AF filtering is then calculated. Next, we determined the number of the detected *de novo* mutations under the same AF criterion. Then, in each Q binning window, the total *de novo* mutations are readily calculated by normalizing the number of the detected *de novo* mutations by the detection rate of the germline mutations. Following the same procedure, we estimated that the levels of *de novo* mutations in three single cells from the expanded clone are 985, 1012 and 1076 respectively (The mean is 1024±47 with the TsTv ratio of 0.96, **Fig. 5C-D**), which is consistent with the true level of *de novo* mutations detected by comparing two bulk sequencing data (Bulk S1 and S2). We also compared the three single cells (S2A-C) to the bulk sequencing data of S2 to estimate the *de novo* mutations introduced during the single-cell expansion, which are 188,188 and 209, respectively (**Fig. S5**). This result shows that the mutation rate during the single-cell expansion is about 7.8 mutation per division. Next, we applied the same filtering criterion and estimated that the average number of *de novo* mutations in the single cortical neurons are 332±92, 226±50 and 273±140 for the three brain samples (**Fig. 5E-F** and **Fig. S6-7**). Our estimated *de novo* mutations are within the range of the recent single-cell expansion-based measurement (200 to 400 *de novo* mutations per cell) ^32^.

Recently, Ochs *et al*. has shown that genome topology plays important roles in safeguarding the genome integrity in response to the double strand breaks ^33^. In this report, we showed that the genome topology also influences the degree of spontaneous DNA damage. Beside the measurement of DNA damage associated variants, we also demonstrated a reliable estimation of the total *de novo* mutations can be achieved for the same single cell. With the ability for simultaneous measurement of dSNVs and *de novo* mutations, LPSSAR offers an unprecedented approach to study genome stability and heterogeneity in various biological and disease systems at single-cell resolution.

## Acknowledgments

We are grateful to the McNair family, Dr. C. Neblett, Dr. Susan Rosenberg, Dr. Arthur Beaudet and Dr. Brendan Lee for their support. We would like to thank Dr. Christophe Herman, Catherine Bradley, Dr. Greg Ira, Dr. Weiwei Dang, Dr. Meng Wang, Dr. Herman Dierick, Dr. Philip Hasty, Dr. Huda Zoghbi and Dr. Brendan Lee for their proofreading and helpful discussion. We would like to acknowledge the Human Genome Sequencing center for their support. We would like to thank Dr. Shawn Zhang for providing the MCF10A cell line. We would also like to acknowledge NIH NeuroBioBank for providing the human brain samples. We would also like to thank other Zong lab members for their assistance during the development of the LPSSAR.

## Author contributions

C.Z. designed the project. M.G. and K.S. contributed to the early development of linear preamplification with double strand conversion strategy. Q.Z. and Y.N. optimized the linear preamplification and developed pipetting-based MDA reaction. Q.Z. and Y.N. performed the LPSSAR experiments of MCF10A and cortical neurons. Y.N. and M.N. performed the single cell expansion experiment of MCF10A. C.Z. and Y.N. performed the data analysis. C.Z. wrote the manuscript. Baylor College of Medicine applied a patent that covers the chemistry of LPSSAR.

## Supplementary Materials

Materials and Methods

Supplementary Fig. 1-8

Table 1-4

