## Supplemental Figures for "Topologically Dependent Abundance of Spontaneous DNA Damage in Single Human Cells"

# Of

### Materials and Methods

#### Cell line and culture

MCF 10A (ATCC CRL-10317) cells were cultured in DMEM/F12 (Invitrogen #11330-032) supplemented with 5% horse serum (Invitrogen #16050-122), 20 ng/mL EGF (PeproTech), 0.5 μg/mL Hydrocortisone (Sigma #H-0888), 100 ng/mL Cholera Toxin (Sigma #C-8052), 10 μg/mL Insulin (Sigma #I-1882), 100 I.U./mL penicillin and 100 μg/mL streptomycin at 37°C, 5% $CO_2$.

#### Human tissues and DNA samples

All human tissues were obtained from the NIH NeuroBioBank under the supervision of the NIH NeuroBioBank guidelines. The three samples were requested from University of Maryland Brain and Tissue Bank. The UMBN are 5554,1740 and 4925. Prefrontal cortex BA8-10 were used.

#### Preparation of neuronal nuclei from Brain tissues

Nuclei were isolated using Krishnaswami et al.'s protocol [1]. Briefly, frozen tissue was first sectioned on ice and immediately transferred to a pre-cooled Dounce homogenizer by 1,500 μl homogenization buffer (250 mM Sucrose, 25 mM KCl, 5 mM $MgCl_2$, 10 mM pH = 8.0 Tris buffer, 1 μM DTT, 1x Halt Protease Inhibitor Cocktail and 0.1% Triton X-100). The tissue section was then homogenized on ice with five strokes of the loose pestle, followed by 12 strokes of the tight pestle. After that, the homogenate was passed through a 40 μM strainer and transferred to a 1.7 mL tube. The isolated neuronal nuclei were then pelleted by centrifugation at 4°C (1000 g, 8 minutes), and resuspended in 500 μl PBS with 0.5% BSA (Fisher BioReagents, 166099A). Blocking of non-specific binding was performed on ice for 15 minutes. 100 μl nuclei were transferred to a new tube for isotype control. The rest of nuclei were incubated with anti-NeuN antibody (Abcam, ab177487) at room temperature for 30 minutes on a tube rotator. To wash the samples, we added 500 μl PBS with 0.5% BSA into samples, followed by centrifugation at 4°C (500 g, 5 minutes). After the removal of supernatant, the nuclei were resuspended in 500 μl PBS with 0.5% BSA and were incubated with Goat anti-rabbit Alexa Fluor 488-conjugated secondary antibody at room temperature for 30 minutes on a tube rotator. Following the incubation, the nuclei were washed with 500 μl PBS with 0.5% BSA and were pelleted by centrifugation at 4°C (500 g, 5 minutes). We then resuspended the nuclei in cold PBS and used Hoechst 33342 for DNA staining. Neuronal nuclei could be identified by green and blue fluorescence under a fluorescent microscopy. Single neuronal nucleus was mouth pipetted into PCR tubes with lysis buffer, and the whole genome was then amplified by LPSSAR. Bulk DNA was extracted using the Quick-gDNA MiniPrep kit.

#### Single cell isolation and alkaline lysis

Cultured cells were dissociated with 0.05% trypsin at 37°C for 15 minutes. The trypsinization was then stopped by growth medium. Following several washes with PBS, single MCF 10A cell was mouth pipetted into a PCR tube containing 2 μL alkaline lysis buffer (400 mM KOH, 100 mM DTT, 2 mM EDTA). After briefly spinning down, the lysis of the single cell is performed by the following temperature: 30°C 1.5 hours. After that, 2 μL (600 mM Tris-HCl, pH=7.5, UV treated), 400 mM HCl) stop solution was added into each PCR tube to neutralize the lysis buffer.

After briefly spinning down, the lysed single cells were ready for the UDG treatment and LPSSAR.

**UDG Enzyme treatment**

UDG Enzyme treatment was performed before the preamplification. 0.2 μL UDG enzyme (New England Biolabs) and 1x ThermoPol Reaction Buffer were first added into the single cell lysate followed by incubation at 37°C for 30 min. The reaction is ready for LPSSAR chemistry. We test on one MCF10A cell without UDG treatment (**Fig. S8**). We observed the significant increase of the variants of C->T/G->A type, indicating the significant number of cytosine deamination were introduced during alkaline lysis. UDG treatment is required to remove this technical artifacts.

**Linearly produced semiamplicon based split amplification reaction (LPSSAR)**

The LPSSAR starts with multiple annealing cycles described below. 300 μM dNTP and 380 nM GAT27NTNG primers are added into each PCR tube containing the lysed cell. In this first cycle, DNA are melted at 94°C for 50 seconds, then the temperature is lowered to 65°C and 2.8 Units of Bst large fragment (NEB) are added into the reaction, after that, transfer PCR tubes immediately to ice to quench the reaction. After quenching for at least 20 seconds, the tube is put back to the PCR block, in which temperature is already lowered to 10°C. Once the tube is placed, we begin the following ramping procedure: 10°C 40 s, 20°C 40 s, 25°C 40 s, 30°C 40 s, 40°C 1 min, 45°C 1 min, 55°C 40 s, 65°C 4 min.

In the second cycle, DNA are denatured at 94°C for 20 seconds, and then at 65°C 2.8 Units of Bst large fragment (NEB) and 0.25 μL GAT27 primer (10 μM) are added into each PCR tubes at 65°C. 8 cycles of 63°C 15 s, 65°C 20 s followed by 1 minute incubation at 65°C are performed to convert all the full amplicons to double strand DNA before quenching. This step will convert the full amplicons to double stranded DNA and avoiding priming to this nonlinearly produced DNA in the next quenching step. The tubes are then quenched on ice and the following temperature steps are performed: 10°C 40 s, 20°C 40 s, 25°C 40 s, 30°C 40 s, 40°C 1 min, 45°C 1 min, 55°C 40 s, 65°C 4 min 30 s (+30 s/cycle). In the third cycle, again DNA are first denatured at 94°C for 20 seconds, at 65°C, 2.8 Units Bst polymerase is added followed by 8 cycles of 63°C 15 s, 65°C 20 s and then 1-minute incubation at 65°C. Then the same quenching procedure is performed.

After the three step of annealing and ramping cycles, 0.2 μL GAT27 primer (10 μM) and $H_2O$ 3.8 μL are added into each tube at 78°C, then DNA are denatured at 94°C for 20 s, and 3.6 Units of Bst large fragment (NEB) are added into each tube at 65°C. Next, 30 cycles of 63°C 15 s and 65°C 20 s were performed and then followed by one step of 2-minute of incubation at 65°C. This extensive DSC step warrants the efficient conversion of all the full amplicons to the double stranded DNA before the MDA reaction. Bst large fragment is inactivated at 72°C for 25 min.

After pre-amplification step, qPCR is performed to quantify the preamplification yield of full amplicons, which is a good indicator of the input of genomic DNA. 10 μL reactions contain 5 μL iTaq Universal SYBR Green Supermix, 0.5 μL GAT27 primers (10 μM), 4 μL $H_2O$ and 0.5 μL preamplification products. The program is as follows: 94°C 2 min for denaturation, then 94°C 20 s, 60°C 25 s, 72°C 2 min 20 s, 28 cycles.

**Semiamplicon based split amplification by MDA**

The reaction was prepared on ice, single cell pre-amplified products was split into 3 tubes (about 4.7 µL each tube) after 10 s vortex at 600 rpm. 24.6 µL MDA master mix including 1x phi29 buffer, 1 mM dNTP, 100 µM random hexamer, 0.5% Tween20, 0.2 mg/ml BSA and about 10 Units of Phi29 DNA polymerase was added into each PCR tube.

After brief pipetting mixing and centrifuging, the tubes were quenched on ice for 3 mins to increase the efficiency of primer binding. Then MDA was performed at 30°C for 25 to 30 minutes with frequent mixing by pipetting. Phi29 DNA polymerase was inactivated at 65°C for 10 mins. After MDA reaction, the products were purified by using 1x AMPure XP beads and eluted into 6.5 µL. The yield was quantified by Qubit High Sensitivity dsDNA kit (Invitrogen Life Science). The yield is around 1 ng of double-strand DNA products per split reaction.

**Library construction**

According to the Nextera DNA sample preparation kit, we carried out tagmentation as follows: 6.3 µL 2X tagment buffer and 0.2 µL tagment DNA enzyme (10-fold dilution using 50% glycerol in 10 mM Tris-Cl, pH 8.5), were added into the 6.2 µL MDA product. The solution was incubated for 1 min 50 seconds at 55°C, followed by adding 1.5 µL 0.2 M EDTA to release transposase at 50°C for 30 mins. Then 1.5 µL 0.2 M $Mg(Ac)_2$ were added to quench EDTA. Then 18.7 µL NEBnext Ultra II Q5 mastermix, 1.5 µL Index 1 primers (N5), and 1.5 µL Index 2 primers (N7) were added. The PCR were then performed with the following cycles: 5 min at 72°C, 30 sec at 98°C, and then 12 cycles of 10 sec at 98°C, 30 sec at 58°C, and 1 min at 72°C, at last, 5 min at 72°C, hold at 10°C. The products were purified by 0.8X AMPure XP beads and eluted to 20 µL Tris-Cl buffer. The libraries are ready for QC as described below.

Qubit High Sensitivity dsDNA kit (Invitrogen Life Science) is used to measure the concentration and the size distribution of the library by the Agilent tapestation. In general, the amplification products have a size distribution of ~300 bp to 1500 bp and the yield is more than 100 ng from each split reaction.

**Loci test and whole genome sequencing**

Before performing whole-genome sequencing, we perform loci test to confirm the amplification evenness. 6 loci were randomly selected from different chromosomes. For each split reaction, when more than one locus drops out, we will discard the cell. Paired end sequencing (150 bp x 2) was performed on a HiSeq X10 instrument with each split have around 10x sequencing depth.

**Bioinformatic pipelines**

The sequencing data of each split is mapped to the GRCh37 (hg19) by bwa algorithm. After the mapping, for the bulk sequencing data, the germline mutations are then called using samtools 1.7. For single-cell sequencing data, GATK3.8 is used for variant calling with the ploidy number set as 10. This way, we can efficiently detect the variants with potentially different amplification bias.

The variants are then compared with the bulk sequencing. To call a variant as a *de novo* variant, two conditions need to be satisfied: first, this variant is not detected in the bulk sequencing data, secondly, there are at least 10 reads in the bulk sequencing data covering this locus (this condition warrants that the *de novo* variant is not false negatives),  Next, we perform a set of filtering using Bedtools to remove the significant tandem regions and the regions that are close to centromere and telomere with high frequency of misalignment. We define as the "vanilla" version of genome. Next, we also remove three bases next to indel to avoid the mis-alignment in these regions. At last, we also implement another filtering script to filter out the homopolymers or the sequence with low complexity. The examples of bash scripts are given online.

**Estimation of the detection rate of dSNVs ($D_{dSNV}$)**

We first derive the strand dropout rate based on the genome coverage of each split MDA reaction: $DR_{strand} = \sqrt[4]{1 - P_{coverage}}$. The stand detection rate is $D_{strand} = 1 - DR_{strand}$. The detection rate of dSNVs base on two-split criterion is estimated as $D_{dSNV} = 3D_{strand}^{2}(1 - D_{strand}) + D_{strand}^{3}$.

**Preparation of single cell expanded clones**

Cells were first serially diluted to 2-5 cells/mL in condition medium. Then, 200 µl of diluted cells were added into each well of 96-well polystyrene tissue culture plate (FALCON). 3 hours after cell seeding, each well was visually checked under a microscopy. Only the wells with one single cell were chosen to generate single cell expanded clones. Single cell expanded clones were cultured for about 25 divisions. The cells were trypsinized with 0.05% Trypsin, single cells were isolated and subjected to LPSSAR, and the rest of cells were used to extract genomic DNA for bulk sequencing using the Quick-gDNA MiniPrep kit.

### Supplementary Figures

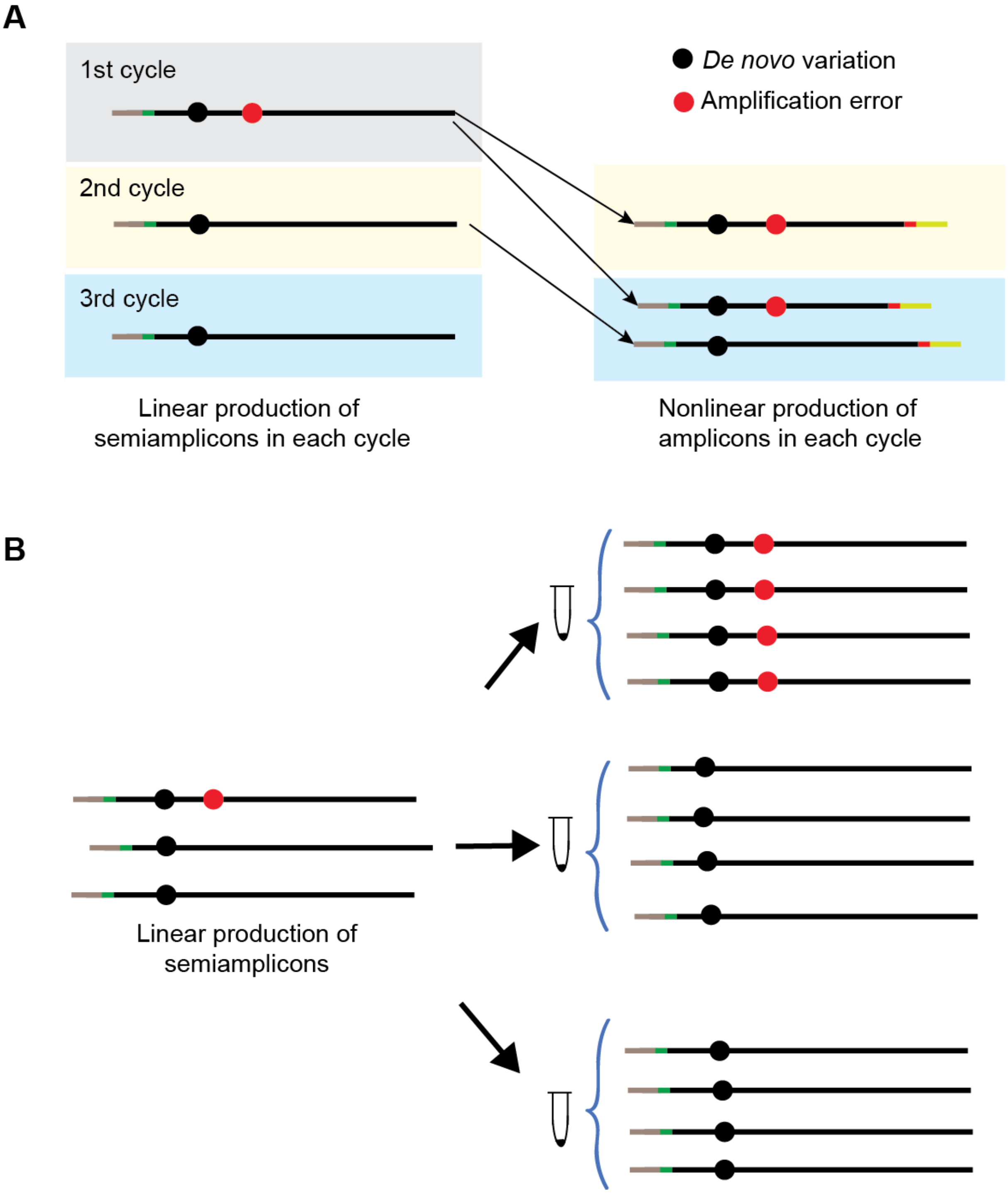


**Supplementary Fig. 1:** (**A**) The linear production of semiamplicons and the nonlinear production of full amplicons in preamplification of LPSSAR. (**B**) The linearly produced semiamplicons are amplified by MDA in each split. The split reactions are required for filtering the amplification errors.

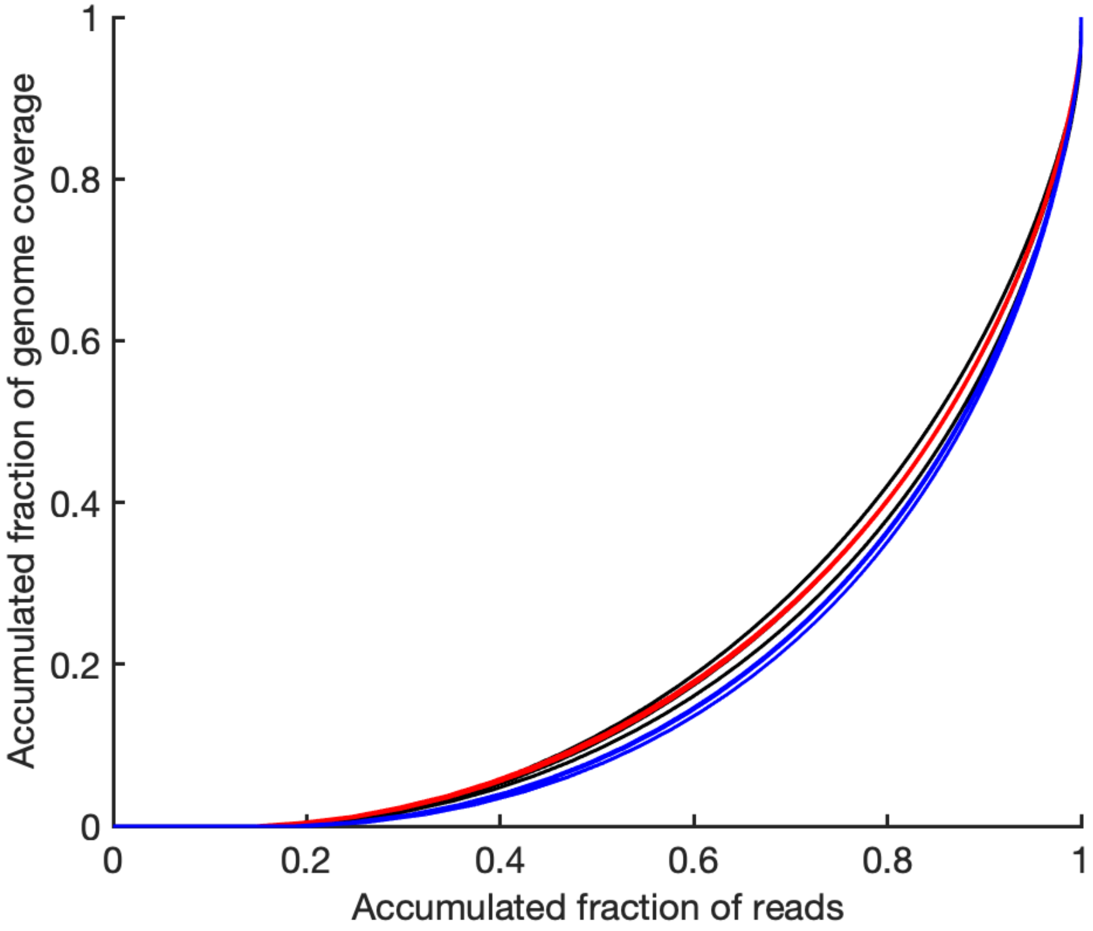


**Supplementary Fig. 2:** Gini plots for the three splits of MCF10A SC1, SC2 and SC3 respectively. The similar curves of the three splits of each cell support the robust MDA reactions were achieved.

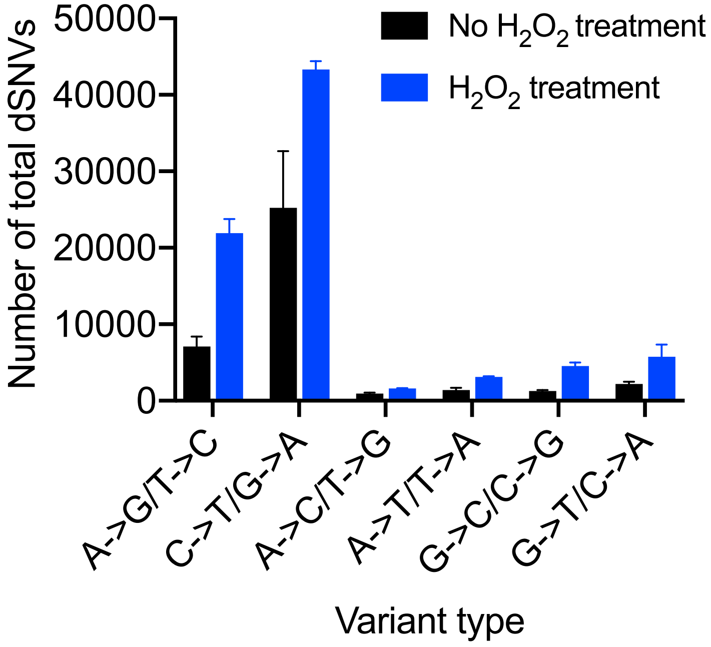


**Supplementary Fig. 3:** The number of different types of dSNVs between the cells with and without hydrogen peroxide treatment in single MCF10A cells.

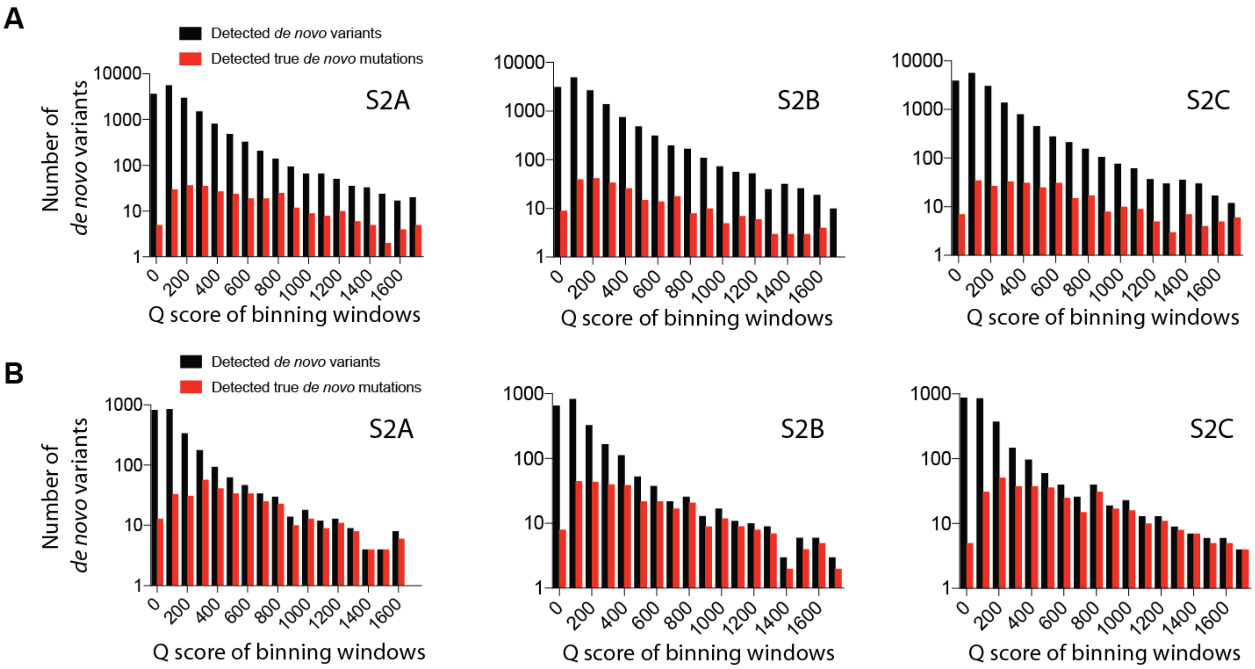


**Supplementary Fig. 4:** (**A**) The numbers of the detected *de novo* transition variants and the true *de novo* transition mutations in the binning windows of the Q scores without the three splits criterion. (**B**) The numbers of the detected *de novo* transversion variants and the true *de novo* transversion mutations in the binning windows of the Q scores without the three splits criterion.

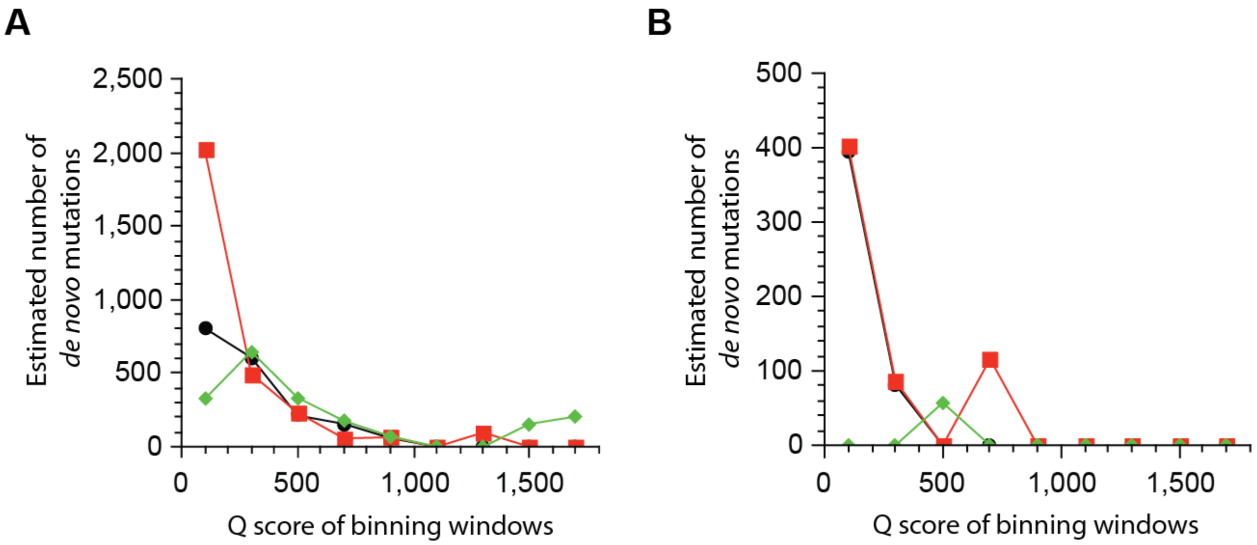


**Supplementary Fig. 5:** (**A**) The estimated total number of *de novo* transition mutations on the axis of the Q score of the binning windows using the stringent AF criterion for the three single MCF10A cells from the single cell expansion (S2A-C). The reference bulk is Bulk S2. (**B**) The estimated total number of *de novo* transversion mutations on the axis of the Q score of the binning windows using the stringent AF criterion for the three single MCF10A cells from the single cell expansion (S2A-C). The reference bulk is Bulk S2.

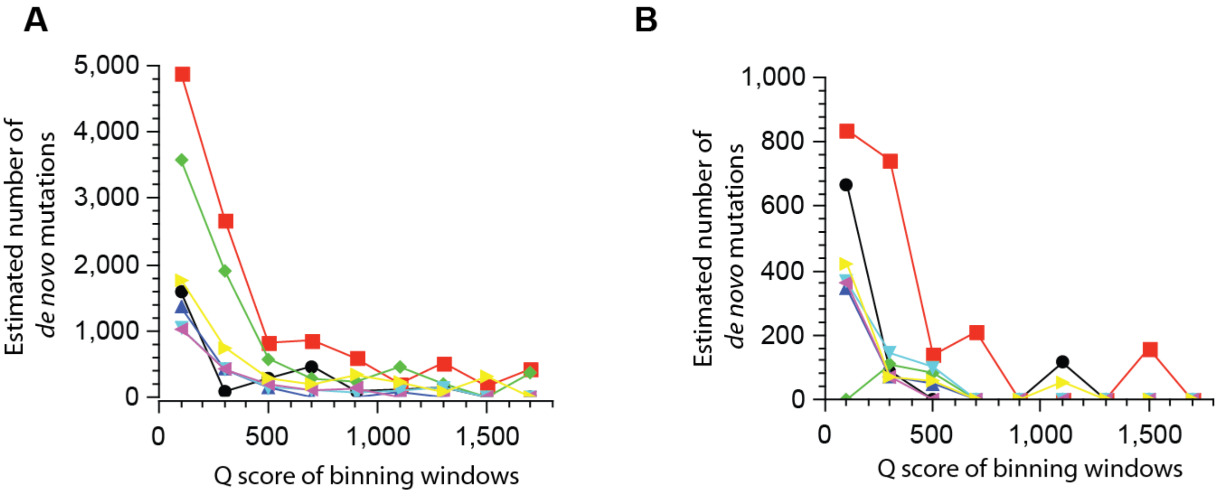


**Supplementary Fig. 6:** (**A**) The estimated total number of *de novo* transition mutations on the axis of the Q scores of the binning windows using the stringent AF criterion for the neurons of the brain sample p1740. (**B**) The estimated total number of *de novo* transversion mutations on the axis of the Q score of the binning windows using the stringent AF criterion for the neurons of the brain sample p1740.

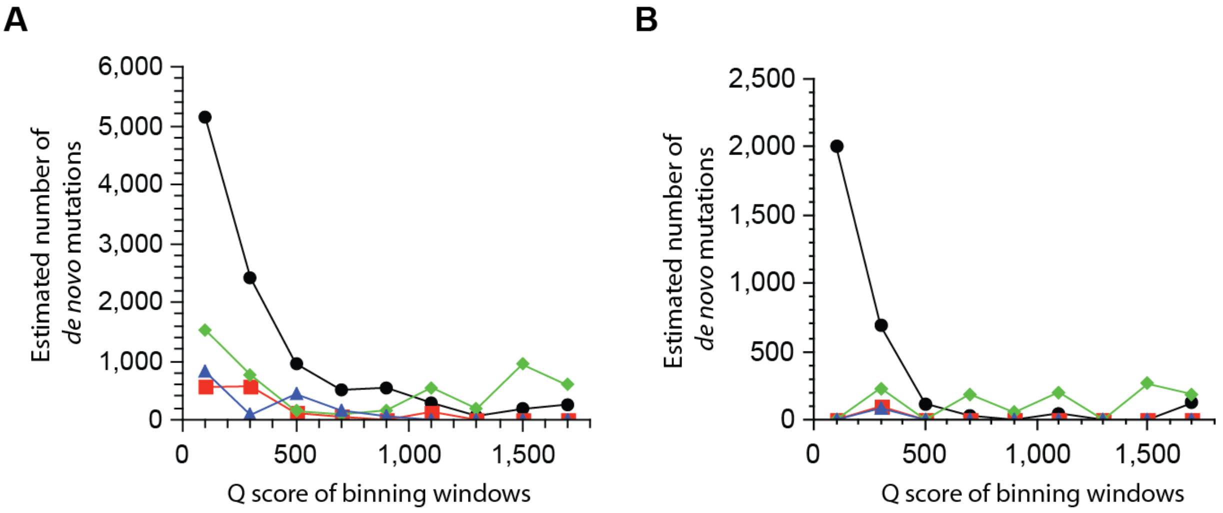


**Supplementary Fig. 7:** The total number of the estimated *de novo* mutations on the axis of the Q score of different binning windows under the stringent AF criterion for the neurons of the brain sample p4925: transition mutations (**A**) and transversion mutations (**B**). For the cell represented by the black curve, only the estimated values at high Q value region (larger than 1500) were used for calculating the *de novo* mutations.

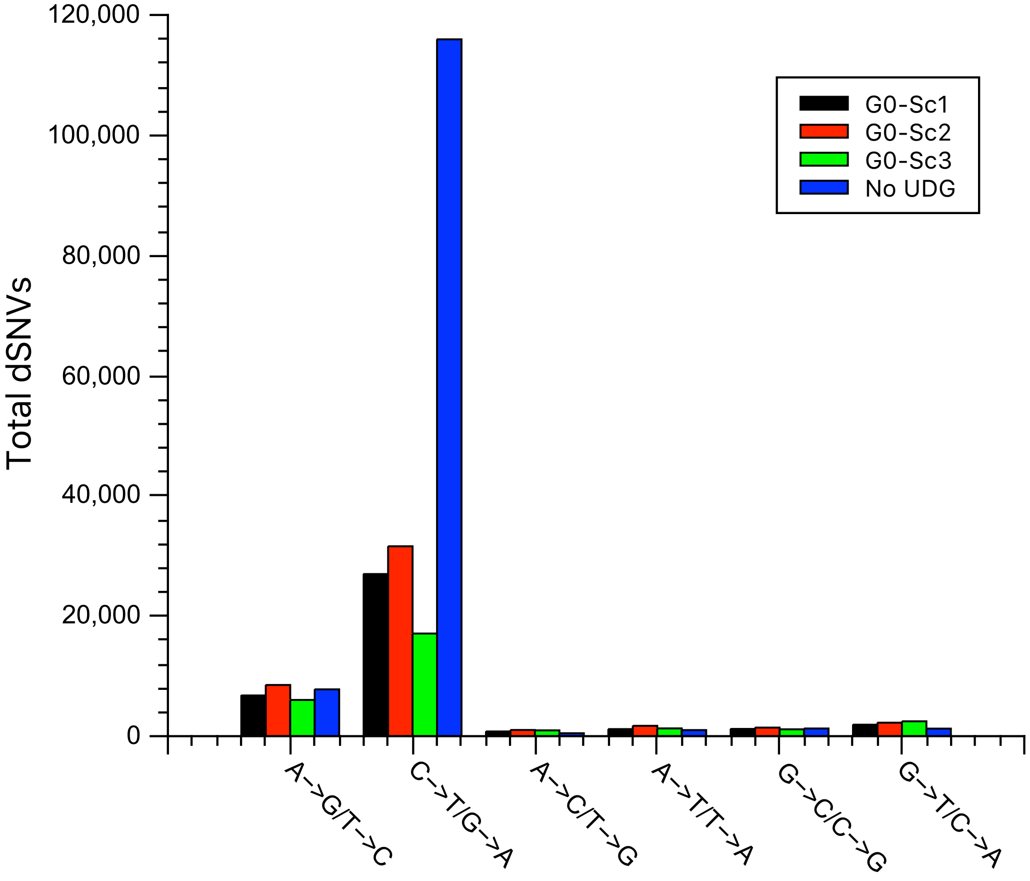


**Supplementary Fig. 8:** The cell without UDG treatment shows significantly higher level of G→A/C→T variants than the UDG-treated cells, indicating successful removal of uracil by UDG treatment.
